# Longitudinal changes in aperiodic and periodic activity in electrophysiological recordings in the first seven months of life

**DOI:** 10.1101/2020.08.18.256016

**Authors:** Natalie Schaworonkow, Bradley Voytek

## Abstract

Neuronal oscillations emerge in early human development. These periodic oscillations are thought to rapidly change in infancy and stabilize during maturity. Given their numerous connections to physiological and cognitive processes, as well as their pathological divergence, understanding the trajectory of oscillatory development is important for understanding healthy human brain development. This understanding is complicated by recent evidence that assessment of periodic neuronal oscillations is confounded by aperiodic neuronal activity, which is an inherent feature of electrophysiological neuronal recordings. Recent cross-sectional evidence shows that this aperiodic signal progressively shifts from childhood through early adulthood, and from early adulthood into later life. None of these studies, however, have been performed in infants, nor have they been examined longitudinally. Here, we analyzed non-invasive EEG data from 22 typically developing infants, across multiple time points, ranging between 38 and 203 days old. We show that the progressive flattening of the EEG power spectrum begins in very early development, continuing through the first several months of life. These results highlight the importance of separating the periodic and aperiodic neuronal signals, because the aperiodic signal can bias measurement of neuronal oscillations. Given the infrequent, bursting nature of oscillations in infants, we recommend the use of quantitative time domain approaches that isolate bursts and uncover changes in waveform properties of oscillatory bursts.

## 1. Introduction

Many drastic changes in brain structure and function occur during the first year of life. Among those are the rapid changes the complexity and diversity of neurons and neuronal connections (Silbereis et al., 2016). In early, prenatal neurodevelopment, these changes are accompanied by a profound shift in cortical electrophysiology, including the spontaneous emergence of neuronal oscillations and their differentiation into multiple frequency bands (Trujillo et al., 2019).

In the context of neuronal oscillations, strong rhythms such as the visual alpha- and sensorimotor mu-rhythms can be readily observed with non-invasive electroencephalography (EEG) in adult humans in the awake state, but these rhythms are not present at birth. These striking differences spurred interest in studying changes of neuronal oscillations across development beginning in the earliest days of EEG (Berger, 1933). Classical cross-sectional and longitudinal studies in infants and children measured oscillatory frequency and amplitude of alpha-rhythms in the time domain, showing that oscillatory frequency increases over the span of childhood, while amplitude decreases (Lindsley, 1938, 1939). The alpha-rhythm emerges at around 3– 4 months, with a frequency of around 3–4 Hz (Smith, 1938; Lindsley, 1939), increasing to 5.5–7 Hz at 12 months; the frequency continues to increase over the course of childhood and adolescence (Henry and Greulich, 1944) before decreasing later in life (Wang and Busse, 1969).

Eventually these time domain approaches gave way to later studies that used spectral domain approaches to measure changes across fixed selected frequency bands in terms of relative or absolute power (Hagne, 1968; Mizuno et al., 1970; Marshall et al., 2002; Saby and Marshall, 2012). However, there are several short-comings associated with using spectral power measures in fixed frequency ranges (Haller et al., 2018; Cole and Voytek, 2019). First, oscillations in infant EEG are transient, appearing in short bursts. This means that spectral measures that average across very long time windows, such as are commonly used, can make very high amplitude oscillatory bursts appear to be much smaller in amplitude, since they average in data from times when no oscillations are present (Jones, 2016; Cole and Voytek, 2019). Because of this, it is unclear whether oscillatory amplitude actually decreases with development, or whether oscillatory bursts become less frequent. Furthermore, an oscillation whose frequency is less stable can also manifest as a lower amplitude rhythm when measured using traditional spectral approaches. Given the differential role that oscillatory bursts play compared to tonic rhythms in neural processing and cognition (Feingold et al., 2015; Lundqvist et al., 2016; Peterson and Voytek, 2017), it is important to clearly elucidate the nature of how these rhythms change with early development.

Another shortcoming of traditional spectral analysis approaches is that EEG activity consists of mixed periodic and aperiodic signals (Haller et al., 2018). Aperiodic activity manifests in the 1/f-like structure of the signal, and is the dominating type of activity when oscillatory bursts are absent. Because of this, even when no oscillation is present, spectral analyses will show power within a frequency band driven entirely by the aperiodic signal, and not by any oscillatory activity. Thus, without explicitly measuring and controlling for the aperiodic signal, one cannot say with certainty whether the band-specific power changes seen in development are driven by changes in oscillatory bursts, the aperiodic signal, or both. This is especially important given the emerging evidence that the aperiodic exponent exhibits strong changes both across aging (up to 70 years old) (Voytek et al., 2015) and across childhood (from 4 to 12 years old) (He et al., 2019). However, these studies are limited by the fact that they are cross-sectional and don’t account for the bursty nature of oscillations, which can also change across development and distort spectral estimates of neuronal oscillations (Cuevas et al., 2014), especially given substantial variability across individuals.

As of yet, no study has explicitly examined longitudinal changes in aperiodic activity and oscillatory bursts across development, which results in a lack of clarity regarding which features are truly changing with development: the aperiodic signal, and/or oscillatory burst amplitude and oscillatory frequency. Therefore, here we aim to show explicit quantification of aperiodic and periodic processes in infant EEG using openly available tools and methodological considerations specific to infant data. For this, we re-analyzed open data from Xiao et al. (2018), consisting of longitudinal EEG measurements of infants in the first seven months of life. We assess changes in the aperiodic component, as well as changes in oscillatory bursts by quantifying waveform features in the time domain (Cole and Voytek, 2019; Schaworonkow and Nikulin, 2019).

We find that the aperiodic exponent exhibits strong changes during infancy, with a marked decrease across the investigated age range of the first month to the seventh month of life. This decrease in spectral exponent is equivalent to a “flattening” of the power spectrum as seen in aging (Voytek et al., 2015), which has previously been linked to an alteration in the relative contributions of excitatory and inhibitory currents contributing to the field potentials from which EEG activity arises (Gao et al., 2017). In addition, we quantified bursts over occipital and sensorimotor regions. While we confirm previous work showing that oscillatory frequency increases with development, we find no significant change in alpha-amplitude. This suggests that prior observations of decreasing alpha-amplitude with development may be at least partially driven by the oscillation frequency changes.

While these results are specific to early development, we argue more generally that the complexity of neural oscillations–including their bursty nature–as well as the often overlooked aperiodic signal, all need to be taken into consideration when assessing spectral measures of neural oscillations. Specifically, we propose an analysis approach that quantifies the aperiodic component and uses burst detection algorithms. Such a combined approach may yield improved clarity regarding the relationship between oscillatory and aperiodic activity to perceptual, cognitive, and motor development in infancy.

## 2. Materials and Methods

### 2.1. Experimental recordings

We reanalyzed an openly available EEG dataset (Xiao et al., 2018; Hooyman et al., 2018). Here, we give a brief summary of the experimental methods, with full details given in the original publications.

#### 2.1.1. Participants

The study protocol was approved by the institutional review board of the University of Southern California. Written consent was obtained prior to the experiment from the parent or legal guardian. The dataset was collected from 22 typically developing infants (10 male, 12 female), stemming from full-term births, with 1–6 recordings per infant and a one month interval between sessions, with the minimum and maximum age present in this dataset of 38 and 203 days, respectively. Participants exclusion criteria were: (1) complications during birth (2) any known visual, orthopedic or neurological impairment (3) below 5^th^ percentile Bayley Scales of Infant Development (Bayley, 2005) score at the age of testing. 71 sessions of approximately 5 minute length were acquired in total. One session was excluded from analysis because only 35 seconds of data was contained in the associated file.

#### 2.1.2. Experimental Design and EEG recording setup

The experimental session consisted of four types of blocks: 1) baseline recording where the infant was presented with a glowing globe toy (40–60 seconds) 2) reaching trial: were a toy was placed in front of the infant, with encouragement to reach for it (20 seconds) 3) non-reaching trial: toy was removed (20 seconds). Reaching and non-reaching trials were repeated five times. 4) baseline recording. Since behavioral annotations were not contained in the dataset, we did not differentiate between conditions and analyzed the recording as a continuous trace. Scalp EEG was recorded from a 32 channel BioSemi active electrode cap (Amsterdam, The Netherlands), with channels arranged in the international 10-20 system. The data was available with a sample rate of 512 Hz.

### 2.2. Data analysis

Data analysis was performed with python using MNE v.0.20.4 (Gramfort et al., 2013), and R (R Core Team, 2017) for calculating linear mixed models using the lme4 library (Bates et al., 2015). The analysis code needed to reproduce the analysis and figures is provided here: http://github.com/nschawor/eeg-infants-exponent.

#### 2.2.1. Preprocessing

First, channels were manually rejected using visual inspection to exclude outlier channels according to excessive noise level and displacement; time segments showing movement artefacts were manually rejected. After that, independent component analysis was performed (FastICA on 2–40 Hz band-pass filtered data, principal component analysis was used as a pre-whitening step to first retain components that explain 95% of the data variance which were subsequently passed on to the independent component algorithm. In one session, the threshold needed to adjusted to 99% to return more than one component, because of very strong noise common to all electrodes). Strong muscle noise and movement artefact components were identified manually with aid of spatial topographies, frequency spectra and component time courses. These components were then projected out (mean and standard deviation of number of rejected components: 1.15 ± 1.13). In general, because the separation between noise and signal ICA components in infant EEG data is less distinct than in adults (Noreika et al., 2020), we rejected components conservatively. The data was re-referenced to a common average reference.

#### 2.2.2. Calculation of aperiodic exponents

The spectral parameterization method and toolbox of Haller et al. (2018) (version 1.0.0) was employed for calculation of aperiodic exponents. In this approach, the power spectrum is modelled as a combination of aperiodic and oscillatory components, which allows distinguishing between oscillatory and aperiodic contributions to the spectrum. Following steps were executed to arrive at the aperiodic exponent measure.

1. For each session, data was split into 10 second segments.
2. For each segment, the power spectrum was computed using the multitaper method (using 1 Hz bandwidth, resulting in 9 discrete prolate spheroidal sequence tapers).
3. The aperiodic exponent was estimated from the power spectrum of each segment. All exponents from model fits satisfying a minimum *R*^2^ value of 0.95 were kept for further analysis.
4. The mean exponent across segments was calculated, obtaining one value per channel for each session.

A challenge in analyzing this dataset is the presence of many artefacts stemming from gross motor movements as well as a high level of muscle noise. As the presence of artefacts influences power spectral estimates, the aperiodic exponent was evaluated across segments of data to increase the stability of the estimate by averaging. The length of segments of 10 seconds was chosen to balance off a long enough segment length for reliable estimation of the power spectrum while obtaining a sufficient number of segments to identify non-stationary outliers.

Settings for the spectral parameterization algorithm were: peak width limits: (0.5, 12.0); maximum number of peaks: 5; minimum peak height: 0.0; peak threshold: 2.0; and aperiodic mode: ‘fixed’. Here, we only take into account the aperiodic component from the power spectrum, discarding estimated peaks.The presence of a high level of muscle noise manifests in the spectral domain as an increased level of >10 Hz power levels. We therefore parameterized the spectra in the frequency range 1–10 Hz to reduce contamination of exponent model fit by an increase in muscle noise.

For statistical evaluation, a linear mixed effects model was fit to the aperiodic exponents with participant as a random effect and age as a fixed effect for each channel independently. We extracted the corresponding t-values and parameter estimates for the fixed effect. Significance was assessed with a hierarchical bootstrapping approach: the exponent values were permuted across sessions within a participant and the first session was placed at an age drawn with replacement from the empirical distribution of age at the first session, with subsequent sessions retaining their spacing. 5,000 bootstrapping iterations were performed, fitting a linear mixed model to the permuted exponent values and generating a null distribution of parameter estimates. p-values were computed as: (1 + number times the true parameter estimate exceeds the null distribution)/(1 + number of bootstrap iterations). Bonferroni p-value adjustment was applied to correct for multiple comparisons across 32 channels.

#### 2.2.3. Calculation of waveform features

The bycycle toolbox (Cole and Voytek, 2019) (version 0.1.3) was used for detecting and quantifying burst features in the time domain. Following steps were executed to arrive at average burst features for each dataset:

1. To extract the signals of interest, we used a Laplacian spatial filter, where from the activity of one center electrode, four surrounding electrodes were subtracted (see inset of Fig. 4A and Fig. 4B for used electrodes).
2. A narrow band-pass filter (finite impulse response filter, 3–7 Hz) was used for identification of zero-crossings.
3. With aid of zero-crossings, cycle features were determined on broad-band filtered (1–45 Hz) data.
4. All cycles that pass predefined criteria regarding amplitude and period consistency as well as relative amplitude extent were classified as bursts.
5. Mean waveform features across burst cycles (voltage amplitude, cycle frequency, peak-trough and rise-decay asymmetry) were computed for each session.

The rationale for using a Laplacian filter is to extract more local signals, less influenced by volume conduction, while maintaining computational simplicity, as recommended by Cuevas et al. (2014). We used three spatial derivations, centered over left and right sensorimotor electrodes as well as over central posterior electrodes. Spatial patterns were calculated with aid of the covariance matrix across channels (Haufe et al., 2014).

**Figure 1:**
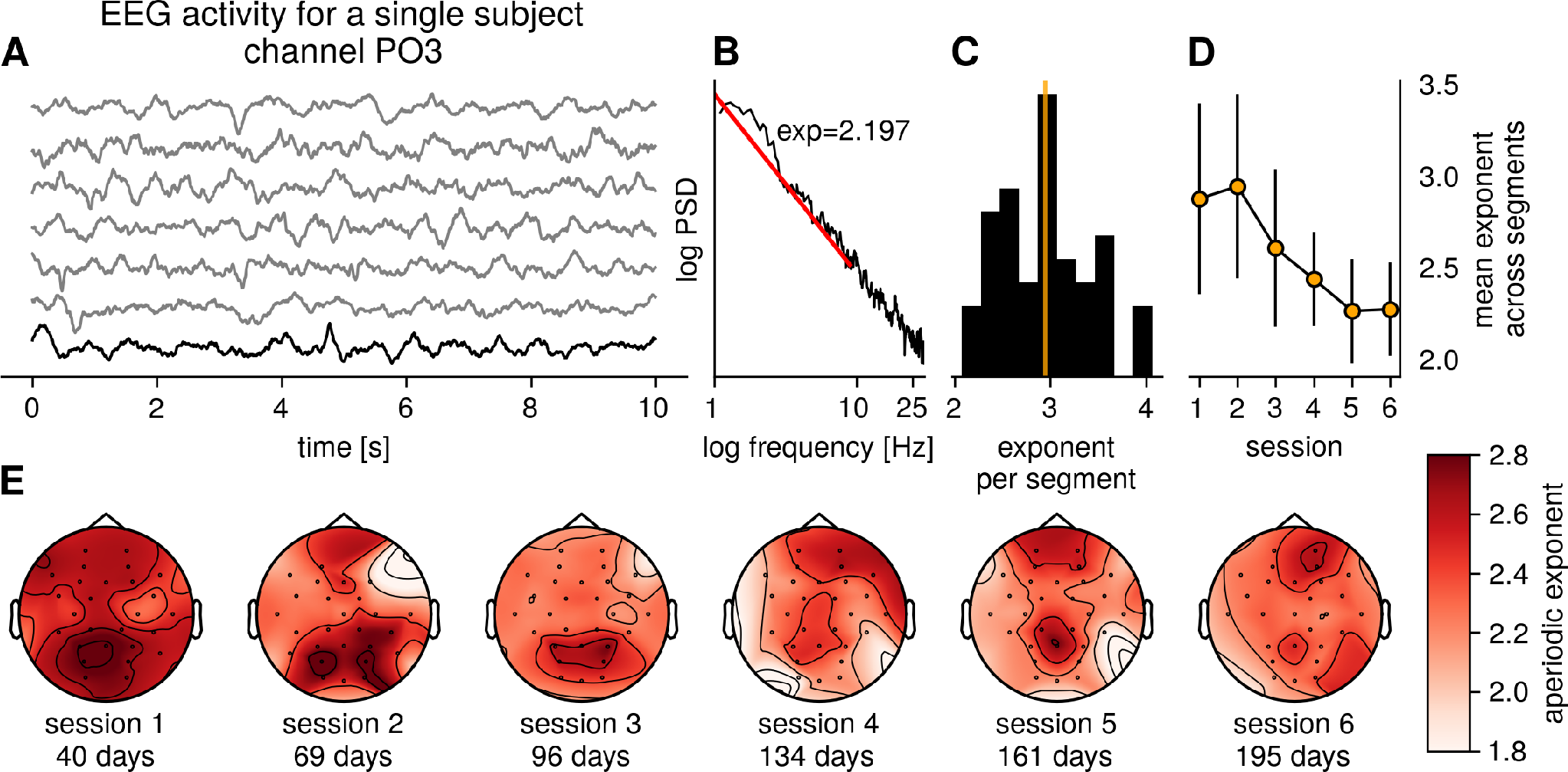
Example exponent calculation procedure and topographies for one subject. A) Infant 32-channel EEG data of approximately was cut into segments of 10 seconds length, with the first six segments shown here. B) The power spectrum was calculated for each channel and each segment, shown is the power spectrum for the black trace. A linear slope was fit to the frequency range 1–10 Hz (red line). C) The aperiodic exponents for segments which had a model fit of *R*^2^ > 0.95 were retained and the mean across segments was calculated (orange line). D) The calculation was performed for each session, error bar shows standard deviation across segments. E) Example topographies of aperiodic exponents for one subject for all sessions, showing gradual decrease in exponent values.

**Figure 2:**
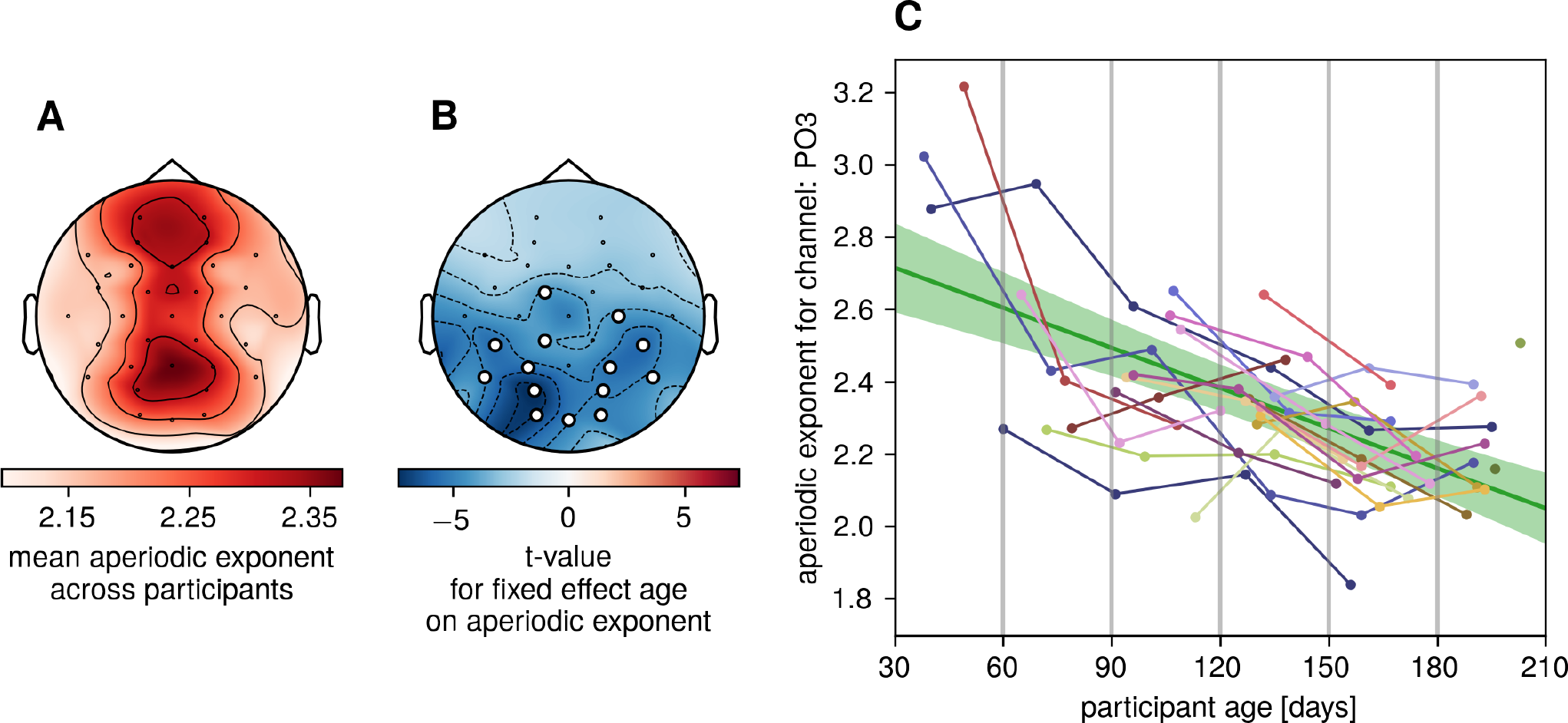
Spectral analysis: aperiodic exponent decreases with age in infants. A) Grand average spatial topography of aperiodic exponent values averaged across all sessions. B) The exponent decreases significantly with age over posterior channels. Shown is the t-value from the linear mixed model for the fixed effect age. Circles mark channels with a corresponding Bonferroni corrected p-value<0.01. N=70 sessions. C) Aperiodic exponent values for the channel PO3. Each line corresponds to one participant. N=70 sessions, Bonferroni corrected p-value = 0.0064. The solid green line is the population level model prediction, with shaded areas representing the 95% confidence interval of the prediction.

**Figure 3:**
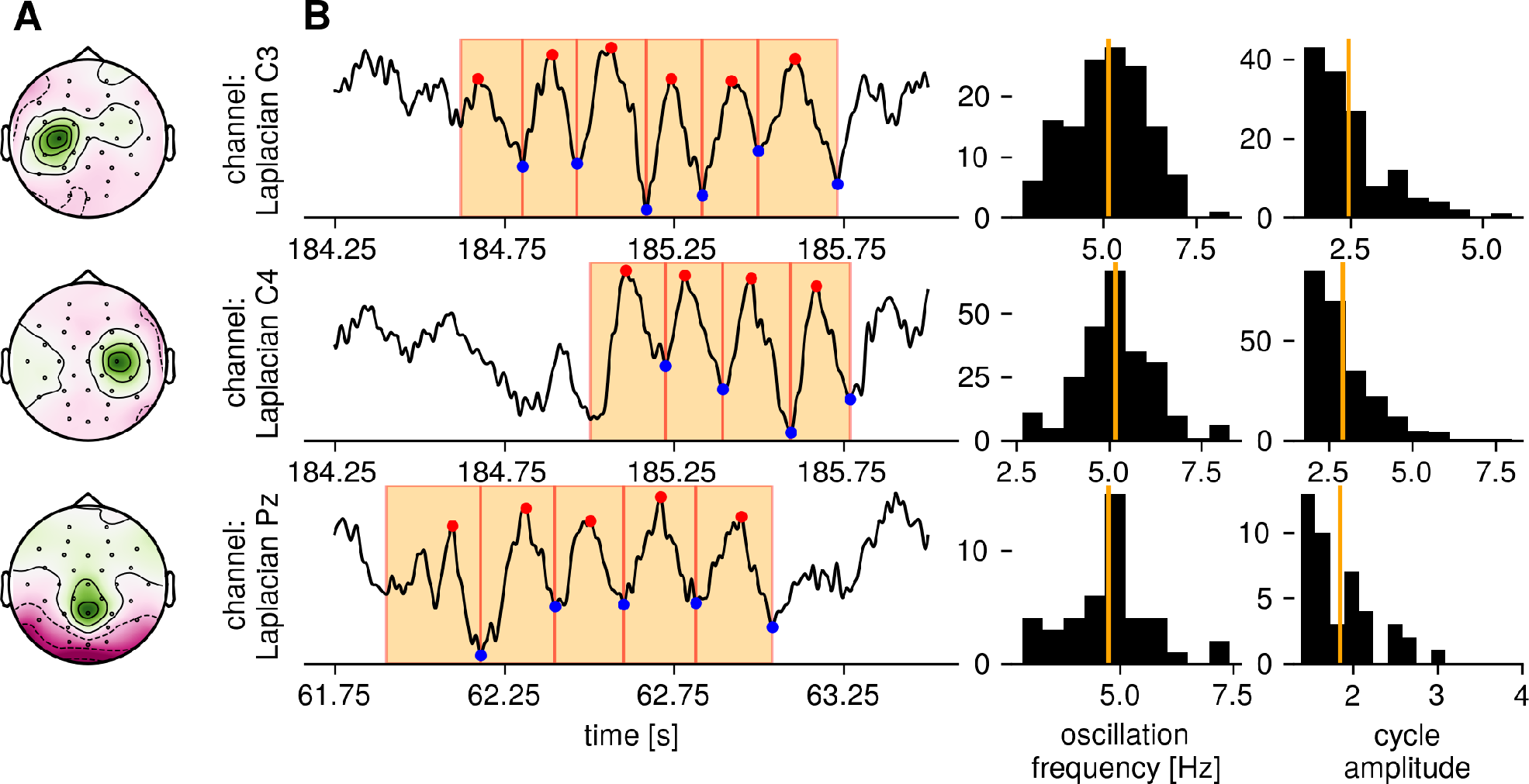
Example burst spatial patterns and time series for one subject. A) Spatial patterns for the three Laplacian filtered channels, which activity was used to perform burst detection, showing a focus over the areas of interest. B) Example traces for each channel with detected burst cycles highlighted. C) After burst detection, cycle features such as oscillation frequency and D) amplitude were computed and then averaged (orange) across each session.

**Figure 4:**
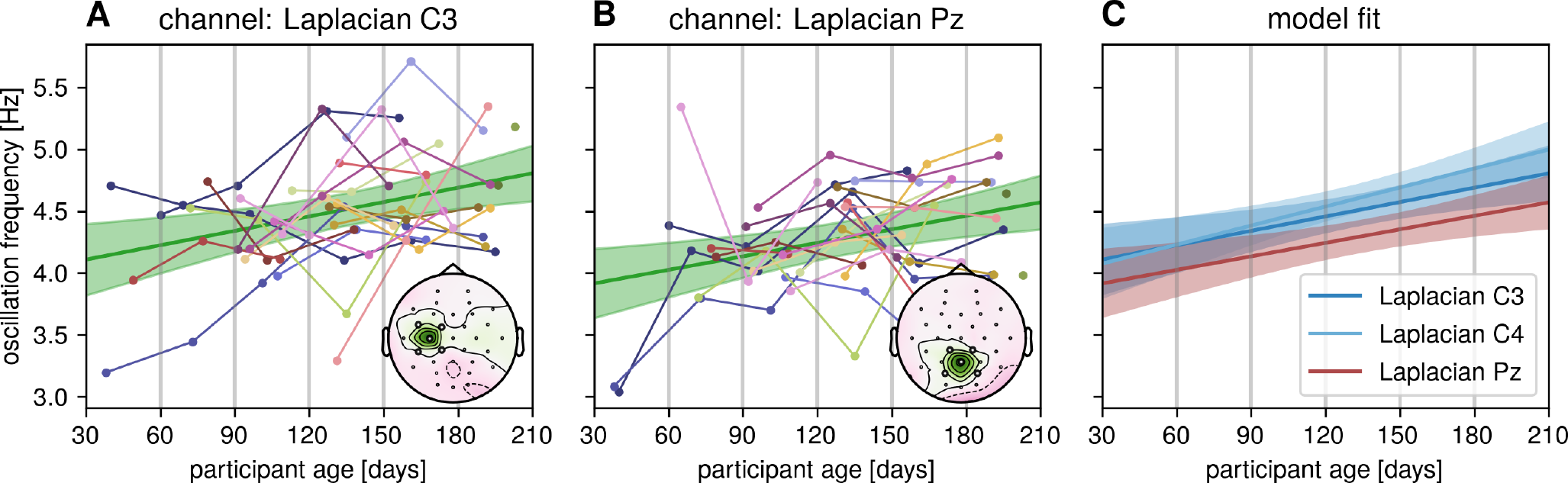
Time domain analysis: oscillation frequency increases with age in infants. A) Oscillation frequency increases with age for sensorimotor bursts. Topography shows spatial pattern for Laplacian C3 channel. N=70 sessions, multiple comparison corrected p-value = 0.0024, obtained by hierarchical bootstrapping. The solid green line is the population level model prediction, with shaded areas representing the 95% confidence interval of the prediction. B) Analog to A), but for posterior bursts as extracted with a Laplacian Pz channel, multiple comparison corrected p-value = 0.009598. C) Mean oscillation frequency for sensorimotor C3 and C4 channels as well as posterior Pz channel increases with age. Solid lines are the population level model predictions, with shaded areas representing the 95% confidence interval of the prediction.

In accordance with previous literature, we chose 3– 7 Hz as the filter range for zero-crossing determination, as this is the frequency range where alpha-rhythm type oscillatory bursts start to gradually emerge in infants. We used the following parameter settings for determining bursts which were consistent across sessions: minimum of three present cycles, amplitude fraction threshold = 0.5, amplitude consistency threshold: 0.5, period consistency threshold: 0.5, monotonicity threshold: 0.5. Even though these criteria are manually selected, they were used for all sessions and all subjects, and thus allow for within subjects comparisons across age.

Analog to the aperiodic component analysis, we then fit a linear mixed model with participant as a random effect and age as the fixed effect separately for each waveform feature and for each of the three Laplacian signals and assessed significance via hierarchical bootstrapping. Bonferroni p-value correction was applied to correct for multiple comparisons across the three Laplacian channels. To compare differences in oscillation frequency across channels, we estimated a common model with data from all three channels with an additional intercept for the factor channel and compared channel-specific intercepts against channel-specific intercepts of a null model (5,000 bootstrap iterations), where waveform-features were pooled and then randomly assigned to a channel.

## 3. Results

### 3.1. Aperiodic exponent decreases with age

The aperiodic exponent was computed for each channel for each session for each participant, the process is illustrated in Fig. 1. The number of analyzed segments per subject (after data cleaning and exluding suboptimal model fits) per session was: 39.54 ± 6.57 (mean ± standard deviation), comprising approximately 94.0% ± 4.3% of the original available data. Example topographies for a single subject across sessions can be seen in the bottom row, showing gradual decrease of aperiodic exponent values. The grand average shown in Fig. 2A shows decreased exponent values for sensorimotor channels in contrast to posterior channels. The exponent decreased with age across the investigated age range for occipital-parietal channels. This was quantified in terms of a negative coefficient of the linear mixed model for the fixed effect age which is shown for each channel in Fig. 2B. The decreasing exponent values across sessions for each participant are shown in Fig. 2C for channel PO3.

### 3.2. Changes across age in oscillatory bursts frequency

We assessed oscillatory bursts in the 3–7 Hz frequency range from Laplacian filtered signals from sensorimotor and occipital regions, see Fig. 3 for example traces for a single subject. The number of detected bursts per session was: 63.50 ± 54.19 (mean ± standard deviation). After burst detection, the mean cycle features were computed for each session and each channel. A linear mixed model showed a significant relationship between mean oscillation frequency and age: oscillation frequency significantly increases with age, for sensorimotor as well as posterior channels (see Fig. 4A and 4B), consistent with previous literature. In this dataset, oscillation frequency of posterior bursts did not differ significantly from frequency of sensorimotor bursts (see Fig. 4C, comparison of intercepts of electrodes of linear mixed model fitted with common slope for age-effect slope, bootstrapped p-value > 0.05). We find no significant effect of burst amplitude and age within the investigated age range, as well as other investigated waveform features (see Table 1).

**Table 1:** Linear mixed model estimates for fixed effect age on average burst features for three Laplacian channels. p-values are obtained by hierarchical bootstrapping. Bonferroni multiple comparison p-value correction across channels and features was applied. N=70 sessions.

| burst feature | C3 – t-value | C3 – p-value | C4 – t-value | C4 – p-value | Pz – t-value | Pz – p-value |
| --- | --- | --- | --- | --- | --- | --- |
| frequency | 3.219631 | 0.0024** | 4.097415 | 0.00240** | 3.176145 | 0.009598** |
| peak-trough asymmetry | 1.084038 | 1.0000 | 0.850820 | 1.00000 | 1.094204 | 1.000000 |
| rise-decay asymmetry | 0.873353 | 1.0000 | -0.449706 | 1.00000 | 0.420949 | 1.000000 |
| amplitude voltage | 0.620464 | 1.0000 | 1.926013 | 0.10078 | -1.672853 | 0.451110 |

## 4. Discussion

The aim of this article is to illustrate robust ways of assessing aperiodic activity in the spectral domain and periodic activity in the time domain for EEG recordings taken during the first months of life. In contrast to adult EEG, sustained oscillations are mostly absent in this data, as indicated by few to no peaks in infant power spectra and corroborated by time-domain analyses, which show that oscillatory bursts occur rarely. We show that aperiodic activity attenuates in the examined developmental interval. The underlying processes that drive this attenuation are not yet clear and need to be studied further in conjunction with structural changes that occur within this time frame, e.g. changes in myelination, brain volume or cortical thickness (Gilmore et al., 2007; Holland et al., 2014), as well as changes in cognitive development. By relating changes in aperiodic exponent to structural and functional measurements it may become possible to distinguish different contributions to the exponent measure, for instance excitation/inhibition balance (Gao et al., 2017) or theories of brain functioning (He, 2014).

Changes in exponent will be reflected in measures of relative power, because normalizing absolute power values will not be sufficient to correct for that. This is because changes in relative power do not necessarily reflect oscillatory dynamics, but can also indicate changes in aperiodic exponent. Care has to be taken to assure that spectral measures used in a study capture the corresponding physiological aspects the researcher intends to measure.

Additionally, we investigated oscillatory bursts in the frequency range of 3–7 Hz using a quantitative method for extracting cycle-by-cycle waveform features for oscillations. This approach allows us to describe changes in waveform features for posterior and sensorimotor bursts. We argue that in the context of oscillation ontogeny, infant data is better suited for time domain analysis as opposed to purely spectral analysis approaches, because of the transient nature of oscillatory bursts. Oscillatory bursts will not necessarily show up in the frequency spectrum computed over long time windows because oscillatory peaks will be eclipsed by the aperiodic component if they occur infrequently (Jones, 2016). Early studies in oscillation changes in development and aging relied on visual inspection and subjective ratings of regularity of rhythms (Lindsley, 1938), but this can be measured by a quantitative assessment of oscillatory bursts, evaluating each cycle in terms of its waveform features.

A limitation of this study is that the functional modulation of oscillatory bursts was not taken into account. While we observed changes in oscillatory frequency with a topographical distribution attributable to visual or sensorimotor rhythms, because of lack in behavior annotation it was not possible to assess rhythm desynchronization with respect to behavior (Stroganova et al., 1999; Bell and Wolfe, 2008), which would be interesting for future studies. Additionally, recordings only had a length of approximately five minutes. During such a short period, the sampled behavior could be very different between sessions, because infant behavior can only be experimentally controlled to a certain degree, making it challenging to assess changes in EEG activity over sessions with this type of data. It would be desirable to run this type of analysis on a larger dataset with longer recordings and behavioral annotations.

## Data Availability Statement

The article is based on openly available data, available in figshare under DOI:10.6084/m9.figshare.5598814 and 10.6084/m9.figshare.6994946.v2. Code to reproduce the analysis is available under: http://github.com/nschawor/eeg-infants-exponent.

